# The BirdsPlus Index, a novel method for assessing site-level conservation values

**DOI:** 10.1101/2025.10.14.682393

**Authors:** Eliot T. Miller, Jeffery L. Larkin, Anna M. Matthews, Michael Parr, Grant Van Horn, James J. Giocomo, Daniel J. Lebbin

## Abstract

While there is growing interest in sustainable management practices to mitigate the biodiversity impacts of agriculture, logging, and other critical societal needs, implementation of such practices is often hindered by a lack of cost-effective, fine-scale metrics that directly link management actions to conservation outcomes. We introduce the BirdsPlus Index (BPI), a novel, scalable approach that integrates monitoring data, remote sensing, and conservation-weighted species scores to quantify deviations of observed site scores from spatiotemporally explicit expectations. Using nearly 29,000 recordings from the Macaulay Library, we generated *acoustic checklists* with the Merlin and BirdNET sound identification models under multiple detection thresholds. We matched these acoustic checklists with species-specific conservation values (BirdsPlus species scores), then trained random forest models to predict total, site-level biodiversity (the sum of these species scores) given environmental and effort covariates. The resulting model also enabled us to map expected BirdsPlus site scores across the landscape.

These scores integrate information on species’ conservation status, ecological roles, and phylogenetic and functional uniqueness. BPI was calculated as the residual between observed and expected site scores, thereby providing a direct site-level measure of conservation value. Across 30 sites, we found that BPI values were consistent across acoustic models and detection thresholds, with high-scoring sites supporting regionally uncommon breeders and habitat specialists. While acoustic- and observer-based (eBird) models showed differing spatial patterns, both aligned with known ecological drivers such as urban density, elevation, and wetland cover. Our results demonstrate that acoustic checklists can be used to model expected biodiversity over time and space, and that the BPI provides a robust, interpretable metric for evaluating the ecological integrity of local sites. Beyond its immediate application to conservation planning, this framework lays the foundation for global, real-time biodiversity monitoring that leverages automated acoustic classifiers, citizen science, and remote sensing to integrate conservation value into development and management decisions.

## INTRODUCTION

Human expansion has been linked to biodiversity loss for millennia^1^, but the strength and consequences of this connection have become increasingly evident in recent decades^2^. Widespread habitat degradation and accelerating climate change have already created the conditions for a growing extinction debt^3^, yet actions taken now could substantially reduce the long-term costs of development to biodiversity^4^. This urgency is amplified by mounting evidence that human wellbeing—financial, physical, and mental—is tightly coupled with functioning ecosystems^5,6^.

As this linkage between human and ecosystem health has grown, so has interest in sustainable development, particularly in approaches such as best management practices for agriculture in biodiverse tropical regions^7^ or conservation incentive programs on working lands^8–10^. While these practices often yield biodiversity benefits and have the potential to affect population trends, their widespread adoption has been hindered by two obstacles: the high cost of documenting conservation outcomes at scale^11^ and the difficulty of financially valuing those benefits^12^. In contrast, short-term declines in crop yields are relatively straightforward to measure^13^; this contradiction can lead to the perception that conservation practices are not financially sound. Addressing this imbalance requires metrics that can both capture the ecological benefits of sustainable practices and link them directly to management decisions.

A wide range of biodiversity metrics has been introduced, but most are either coarse in scale or costly to apply at fine resolution^14^. Map-based indices such as the STAR metric^15^ provide valuable regional insights into the potential gains from restoration or threat abatement, but they rarely offer feedback loops for assessing local-scale actions. Conversely, intensive before–after monitoring projects can detect local impacts, but these are expensive, challenging to scale, and sometimes difficult to interpret. For example, local population stability of a threatened species may actually indicate conservation success in the face of regional decline. Such approaches also frequently overlook the importance of functional traits and the ecological roles of the species in question. Thus, existing approaches leave a critical gap: the absence of scalable, fine-resolution, ecologically informed biodiversity metrics that are cost-effective and directly tied to land-use and management outcomes.

Here we propose the BirdsPlus Index (BPI), a biodiversity index designed to provide near real-time, spatiotemporally explicit assessments of conservation outcomes. The BPI leverages acoustic monitoring, which combines cost-effectiveness and scalability with the capability to standardize data collection across sites. Moreover, because recordings are archived, detections can be both manually verified or revisited later, as automated sound classifiers improve over time. By relying on such classifiers, which have quantifiable error rates^16,17^, this approach avoids a major limitation of traditional surveys: variation in observer skill. While acoustic methods face challenges—including false positives and incomplete species coverage—these issues are surmountable. False positives are a pervasive but often underappreciated problem in biodiversity monitoring generally, with known methods of mitigation^18^, and acoustic classifier coverage and accuracy continue to expand rapidly^19^. To diminish errors, we apply species-specific detection thresholds derived from a vetted test dataset, while also exploring a range of flat thresholds. For demonstration, we focus on the Northeast United States, where, owing to a legacy of focal recordings of known identity^20^, species coverage in current classifiers is already sufficient.

Birds are an ideal taxon for this effort. They are taxonomically and ecologically well studied, with extensive information on distributions, traits, and ecological roles^21^. Their diagnostic vocalizations make them comparatively easy to detect acoustically, and they are especially well represented in current automated classifiers^22,23^. By centering the BPI on birds, we can move beyond species richness alone to incorporate ecological traits, population trends, and habitat preferences into biodiversity and conservation assessments.

Beyond its immediate application, the BPI establishes a framework for scaling acoustic biodiversity monitoring globally and integrating with established initiatives such as eBird^24,25^. Standardized^26^ and semi-structured monitoring approaches like eBird are particularly well-suited to quantifying population status and trends because they include information not only on what was detected, but what was not^27,28^; detections are submitted in the form of complete *observer checklists*. Such approaches are complicated, however, by the participation of numerous observers with varying skill levels^29^, and strong and varying^30,31^ spatiotemporal biases in observer effort. The current challenges are well understood, fortunately^25,32^, and can largely be accounted for, but new complications regularly emerge. For example, mobile phones equipped with apps for automated sound identification (like Merlin) have been widely adopted by the birding community over the past five years, and these tools are almost certainly having detectable impacts on reporting rates and downstream biodiversity inferences. Passing sound files through acoustic classifiers like BirdNET^22^, and applying thresholds to the resulting predictions creates what we define here as an automated *acoustic checklist*. Acoustic checklists generated from classifier outputs parallel observer checklists by recording both detections and non-detections, but with distinctive advantages: error rates can be directly estimated, sound files can be re-analyzed as classifiers improve, and the methods are accessible to a wide pool of participants, from field biologists to casual users of mobile bird identification apps. While automated acoustic monitoring cannot replace direct observation, it provides a scalable and standardized complement to existing monitoring programs^33^. Most importantly, acoustic approaches allow direct comparisons between observed biodiversity and modeled expectations, enabling robust assessments of management and conservation outcomes.

The BPI is powered by four key data resources: (1) BirdsPlus species scores^34^, which integrate functional^35^, phylogenetic^36^, spatial^37^, and population-trend datasets^27,38^; (2) acoustic classifier predictions from BirdNET^22^ and Merlin; (3) the eBird database and Macaulay Library, which collectively house billions of bird observations and millions of sound files^24,25^; and (4) remotely sensed spatial data derived from satellites^25^. These complementary resources provide the spatiotemporal foundation for robust biodiversity and conservation value modeling.

In this study, we test the hypothesis that the conservation value and ecological integrity of sites can be better understood by processing large volumes of acoustic files with a single automated classifier and linking predictions with habitat and sampling effort variables to build spatiotemporally explicit models. We focus on the Northeast United States, a region with strong data density and species coverage, and use the BPI to evaluate the prediction that small, well-sampled greenspaces exceed regional conservation value expectations. In parallel, we generate an analogous set of results using only eBird data, allowing us to compare automated acoustic monitoring with traditional observer-based approaches and highlight their potential integration.

## METHODS

### Data sources

Our study used the recently released BirdsPlus species scores^34^ as our source of integrated conservation assessments for this project. These species-level scores theoretically range from 0-3, although scores for the species included in this study ranged from 0.79 to 1.78 with higher values indicating species of greater conservation value. Community-level samples of these scores tend to be normally distributed^34^. To emphasize the significance of species of conservation concern, we cubed these raw BirdsPlus scores for downstream analyses. The resulting scores ranged from 0.49 (American Robin, *Turdus migratorius*) to 5.66 (Black Rail, *Laterallus jamaicensis*).

We employed a global data resource of increasing prominence and importance: the eBird enterprise writ large, which includes not only the single largest source of biodiversity data^39^, but also the largest curated natural history multimedia library in the world, the Macaulay Library. It is from the Macaulay Library, and its growing archive of three million sound files, that we drew from to create the acoustic checklists we used to generate our expectations, and it is also widely used for building and testing acoustic classifiers. We define an acoustic checklist as a list of all species detected, with high confidence, across a given audio recording.

Our environmental data came from a variety of freely available satellite and other remote sensing spatial layers (elevation, habitat type, etc.). We used 137 such layers in total, and provide the full list with citations of layers in Table S1. These are the same inputs used in the eBird status and trends models, and they are summarized over a 3km diameter circular neighborhood around each checklist location, taking the mean and standard deviation for continuous features, and percent cover and edge density for categorical features^27^. For mapping purposes, we predicted our spatiotemporal model of expected BirdsPlus site scores to a surface of the environmental layers, and we fixed our effort variables (date, time, recording length; see below) for such maps to the values that maximized the predicted site-level BirdsPlus scores.

When an eBirder submits a checklist of birds detected, they can choose to include photographs or audio recordings of those species, which are then stored and managed by the Macaulay Library. Our acoustic checklists are based on every recording (n = 28,798) from a complete eBird checklist (n = 15,055) between 2008 and 2022, with an associated recording in the Macaulay Library in the states of New York, Pennsylvania, Ohio, West Virginia, Virginia or Maryland from the months of June, July or August. These months roughly correspond to the breeding season in these states. As described elsewhere^25^, a complete eBird checklist is the result of a single sampling (bird watching) event where the observer indicates that all detected birds were reported. This allows inference to be made about non-detection. It bears emphasizing that by focusing only on complete checklists with associated audio files—useful for directly comparing these detection modalities—we likely shifted the distribution of eBird observers away from the mean (e.g., on average our dataset might include more skilled observers), and we certainly limited the size of both the eBird and acoustic datasets. For example, there are many more complete eBird checklists in the region that do not contain audio files, and there are many incomplete eBird checklists that do contain audio files. In our dataset, there was an average of 1.91 recordings per observer checklist. The average recording length, which is the acoustic checklist equivalent to the duration of an eBird observer checklist was 78.53s (SD = 123.40). The average eBird checklist length in our sample was 6519.28s (SD = 4712.96). A schematic of our full workflow can be seen in Fig. 1.

**Figure 1.**
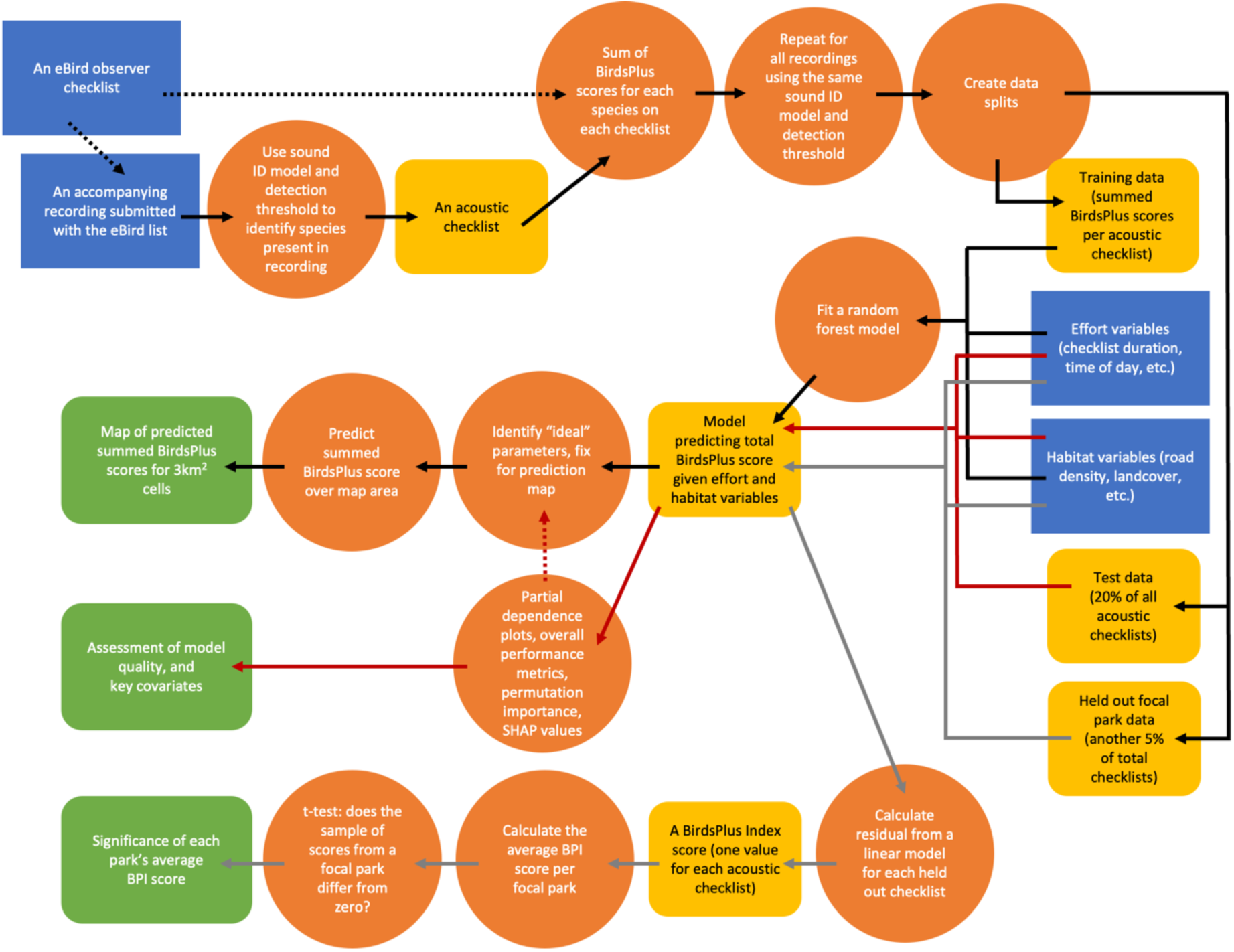
Schematic diagram of the analytical workflow used in this study. Blue rectangles show data sources. The diagram is structured around the workflow specific to acoustic checklists, but the workflow is nearly identical for eBird observer checklists, and the dashed black arrow at the top of the figure indicates this. Orange circles show methodological steps, while rounded, yellow rectangles show intermediate data products. After splitting the data into train, test, and focal park datasets, each split is combined with effort and habitat variables and passed through a separate stage of the workflow. This is indicated by the black, red, and gray arrows, which culminate in three rounded, green rectangles that show the final outputs of the workflow.

### Creating acoustic checklists

We used BirdNET (v2.4) and the Merlin (v43) sound ID models to derive acoustic checklists. Both of these operate by taking as input a 3s-long segment of audio, which is then converted to a spectrogram (a visual representation of sound) and passed through a multilabel computer vision (AI) model. By default, Merlin has a stride of 1s, such that every second of audio is analyzed three times (and there is an output for every second of audio); we modified the BirdNET defaults (by setting “chunk_overlap_s” to 2) to follow this same approach. Thus, the final output from each model is a continuous “probability of occurrence”, from 0-1, for every species in the model (1,789 in Merlin, ∼3,000 in BirdNET) for every second of a recording. These values are not true probabilities of occurrence, however. In general, larger values correspond to higher confidence that a species is truly present in the 3s clip, but there is an imperfect relationship between true presence and these values (e.g., some large prediction scores will still be false). Moreover, if we were to model this relationship as a logistic function, then the inflection point of the curve would vary widely across species^40^. For Merlin, we were able to leverage results from a held-out test dataset to make reasonable estimates for species-specific detection thresholds to implement.

Specifically, we selected thresholds that, on a per-species basis, were estimated to meet a 95% precision criterion. However, because we did not have access to a comparable held-out test dataset for BirdNET, we also set single-value (flat) detection thresholds for both sound ID models. We employed three such thresholds: 0.25, 0.5, and 0.9, which resulted in seven complete sets of acoustic checklists. Per model, per threshold, therefore, a species was considered present on an acoustic checklist if it was detected at least once above the relevant threshold in the corresponding sound file (subject to the following spatiotemporal filtering steps).

BirdNET and Merlin are both global models and, by default, outputs are not spatiotemporally constrained. To reduce the impact of potential false positives, we filtered results by defining a list of acceptable species and zeroing out predictions for disallowed species. This list was defined, per acoustic checklist, by referencing the corresponding eBird dataset— allowable species were those detected by an eBirder within a two-week period and a 3-decimal-degree grid cell centered on the acoustic checklist in question.

### Assembling the training and test datasets

To derive a site-level measure of conservation value, we joined the species on each observer and acoustic checklist with their corresponding cubed BirdsPlus scores, then took the sum of these scores per recording. Next, we matched each sampling event to environmental data as described above. Last, we matched each checklist to a set of sampling effort variables. Common across both detection modalities were: the duration of the checklist itself, the year, day of the year, time of day, time relative to solar noon, distance traveled (which was set to zero for acoustic checklists), and the number of observers on the eBird checklist. To observer checklists we obtained and added an additional covariate: checklist calibration index (CCI), a measure of observer expertise^29^. To acoustic checklists we calculated and added two additional covariates: the sum of acoustic complexity index^41^ values per bin in the sound file, and the entropy of the average spectrum^42^, both calculated using the default parameters in scikit-maad^43^. Because it seems likely that the ability to correctly identify species is correlated with signal to noise ratios, we included these covariates as a way of calculating how “noisy” a recording was.

To evaluate the performance of our models (see below), we created test datasets by setting aside 20% of the data (in addition to another 5% for the case study described below). Rather than randomly selecting test points, which would have perpetuated known spatiotemporal biases in the eBird dataset^25^, we created a spatiotemporally stratified subsample as follows. First, we overlaid a 100x100 grid of cells over the study area, then identified the cell and year to which each acoustic checklist belonged. In our study, that corresponded to cell sizes of 0.08° latitude and 0.13° longitude. Second, we found the total number of checklists in each cell-year combination. Third, we took the reciprocals of these cell-year totals, and used these as a probability of randomly sampling a checklist for inclusion in the test dataset. This has the effect of increasing the likelihood that a checklist from a poorly represented cell-year combination is included in the test dataset.

### Assembling a case example

Because the goal of our study was to develop an index that can be used to assess the relative conservation value of a specific site, we wanted to explore its performance in an applicable setting. As described above, we therefore focused on small, well-sampled parks in the study area, and predicted that such parks, when situated in otherwise large urban regions, would support biodiversity of higher conservation value than expected for that time and place. We were motivated to focus on well-sampled parks, particularly given our generally short-length recordings, because a large number of samples is desirable before conclusions are drawn about the conservation value of the site. Accordingly, we focused on parks from which at least 20 recordings had been made during the study period (June-August). We chose to focus on small parks due to the scale of our environmental data. Specifically, because we summarized our environmental data in a 3km diameter circle around each acoustic checklist, predictions from our fitted model correspond to expectations for an area of 7.07 km^2^. As such, we focused on parks that were, at the maximum, half that area such that site-level expectations were informed by a broader habitat context.

To identify which park, if any, a recording came from, we downloaded the Complete U.S. ParkServe dataset from the Trust for Public Land (https://www.tpl.org/) on 9 December 2024, and intersected the location of the acoustic checklists with the park boundaries. We calculated the area of each park using the *sf* package in R. Thirty parks met our data thresholds for inclusion. The number of recordings per park ranged from 21-147, and the size of the parks ranged from 0.16-3.51 km^2^. Similarly to the test dataset, the summed, site-level BirdsPlus scores from these 1,330 recordings total were withheld from training.

### Model training

Once our training data were assembled—summed, site-level BirdsPlus scores from each of the eight complete sets of checklists (three BirdNET detection thresholds, four Merlin detection thresholds, and eBird observer lists), plus associated environmental, sampling effort, and other covariates—we used a random forest model to predict a site-level BirdsPlus score (Fig. 1). We log-transformed the summed BirdsPlus scores before fitting the model, then exponentiated the predicted scores back to the original scale for downstream use (because we included checklists with no species detected in the model, we used the NumPy^44^ function log1p, and its inverse, expm1 when transforming the data). This transformation was done to combat underprediction of large total BirdsPlus scores. We also explored XGBoost model fitting^45^ for similar reasons, but found it offered no improvements over the random forest model.

We fit the random forest regression model using the scikit-learn package (v1.5.2) in Python. To optimize model performance, we conducted a randomized search over a hyperparameter grid. Specifically, we used the RandomizedSearchCV function to evaluate combinations of the number of estimators (n_estimators = 100, 300, 500), maximum tree depth (max_depth = 10, 20, or unlimited), minimum samples required to split a node (min_samples_split = 2, 5, 10), minimum samples required at a leaf node (min_samples_leaf = 1, 2, 4), and the number of features considered at each split (max_features = ‘sqrt’, ‘log2’, or all).

The randomized search tested 20 parameter combinations using 3-fold cross-validation, and model performance was evaluated based on coefficient of determination (R²). We used the best-performing model from the hyperparameter search for further analysis.

### Model testing

We tested model performance using a number of approaches. First, we used partial dependence plots to assess the relationship of the effort and acoustic complexity variables to the modeled, expected BirdsPlus site scores. In the process, we found the values of these covariates that maximized the expected score and retained those for mapping purposes (see below, also Fig. S1). Second, we used the effort metadata and habitat information associated with each test point to predict its true total BirdsPlus score. We exponentiated the logged scores to convert them to the original scale before calculating the following overall performance metrics: root mean squared error, mean absolute error, and R^2^ of the predicted versus the actual scores. Third, we employed the permutation importance function in the scikit-learn package to quantify feature importance^46^. Fourth, we used the Python shap package to calculate and visualize SHAP values, an “explainable AI” method of understanding feature importance^47^.

### Spatial biodiversity expectations

We visualized our model predictions by using a spatial prediction grid with environmental values averaged across 3km grid cells, and where the values that maximized the additional covariates were appended to each grid cell (effort, acoustic complexity index, etc.). Thus, for example, in the map shown for Merlin with a flat threshold of 0.9 (Fig. 2), the day of the year is fixed at June 1, time of day at 5:42 AM, and checklist length to 170.4s.

**Figure 2.**
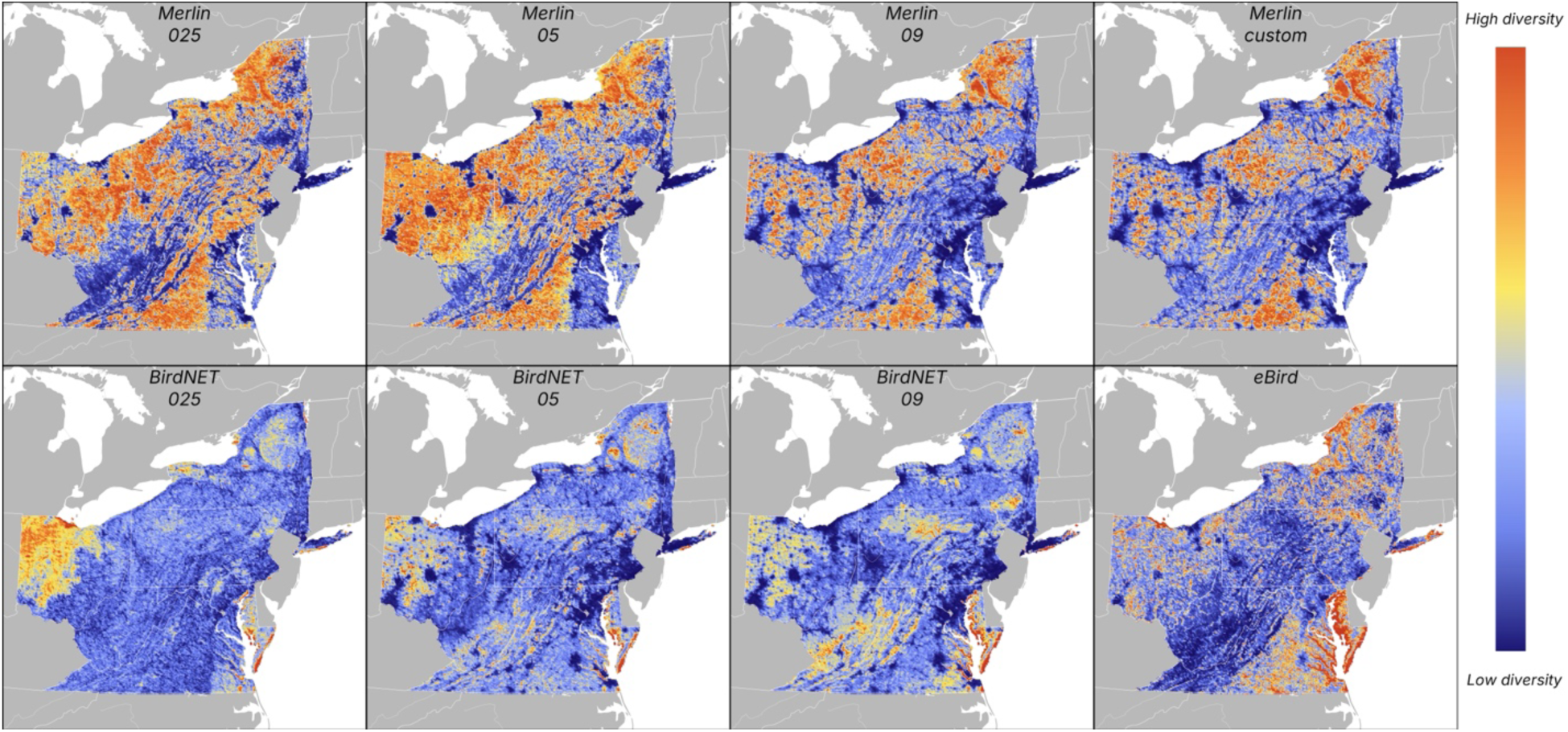
Maps of expected total BirdsPlus scores show where high and low levels of avian biodiversity are expected across northeast USA. These maps indicate expectations during optimal detection conditions for each detection modality and threshold.

Detections from eBird observer and acoustic checklists are fundamentally different, and integrating these detection modalities requires additional research. Nevertheless, to determine how predictions from each acoustic-based model compared to those from the observer-based model, and to other acoustic models, we also retained the pixel-level model predictions from each map and calculated the correlations amongst these.

### Deriving the BirdsPlus Index

We derived the BirdsPlus Index (BPI), a relative measure of conservation value as compared to spatiotemporally calibrated expectations, as the residual, per checklist, of a linear model of the observed site-level BirdsPlus scores as a function of their predicted score (Fig. 3). Thus, checklists with large positive residuals correspond to those where higher BirdsPlus site scores were detected than expected for that time, location, and degree of sampling effort. We repeated the process of calculating observed BirdsPlus site scores and comparing them with expectations eight times (seven acoustic models/detection thresholds plus one eBird model).

**Figure 3.**
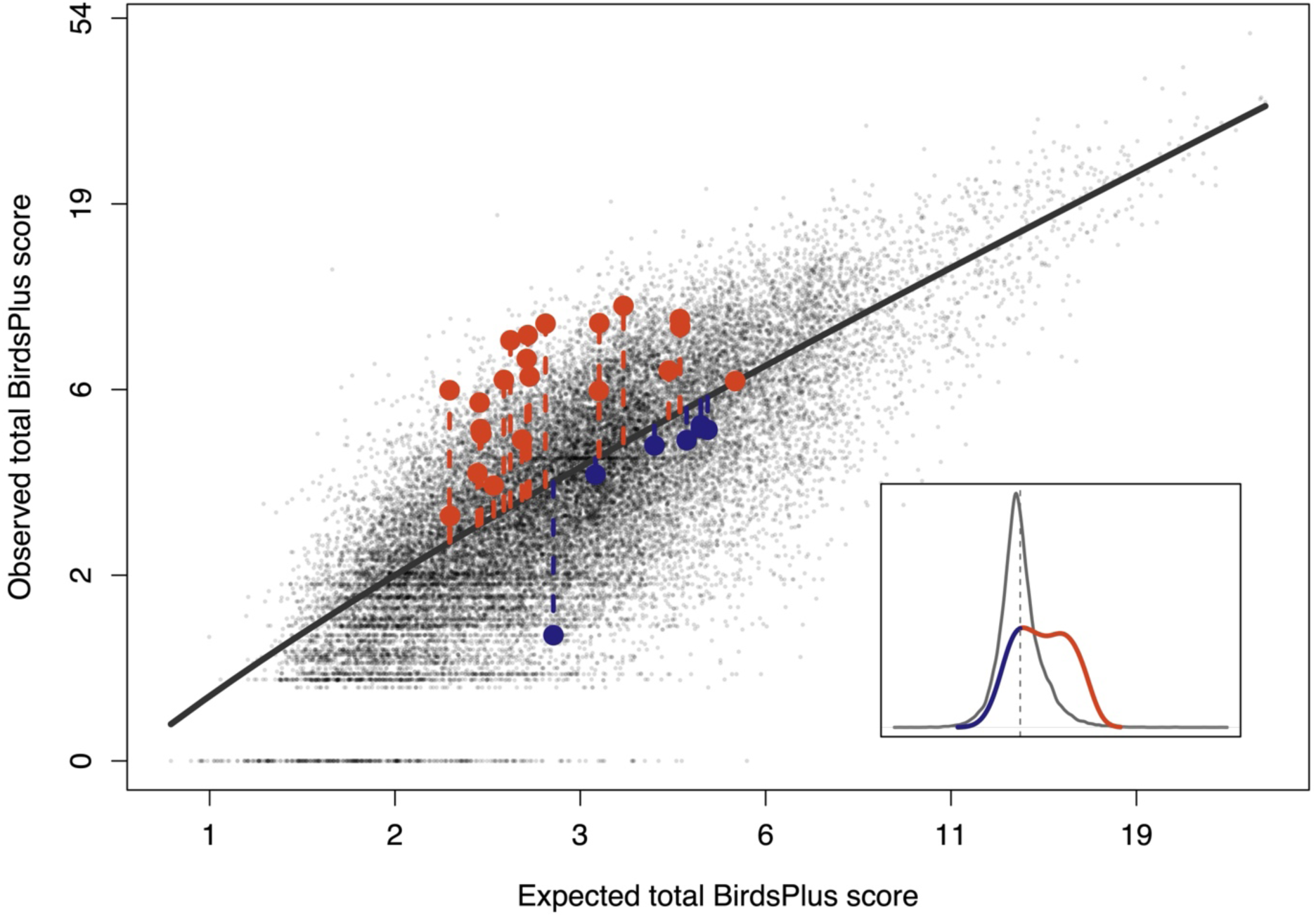
Plot of observed versus expected total BirdsPlus scores per acoustic checklist for Merlin with species-specific detection thresholds show the reasonable performance of the model overall (R^2^ for test dataset = 0.42), but also the systematic underprediction of total BirdsPlus score in the highest scoring sites. The orange and blue points show acoustic checklists from Pennsylvania State Game Lands 117, a park with significantly larger BirdsPlus Index scores than expected from the regional model. The park hosts breeding populations of locally or regionally uncommon species including Henslow’s Sparrow (*Centronyx henslowii*, BirdsPlus species score = 1.31), Yellow-breasted Chat (*Icteria virens,* BirdsPlus species score = 1.04), and Prairie Warbler (*Setophaga discolor*, BirdsPlus species score = 1.27). The inset density plot shows the residuals (BirdsPlus Index scores) for all acoustic checklists in gray, as well as the distribution of residuals for acoustic checklists specific to Pennsylvania State Game Lands 117.

We calculated BPI scores for each of the 1,330 recordings (or 546 eBird checklists for the eBird model) from the 30 parks that met the data thresholds. Per park, we then calculated a mean BPI score to represent its average conservation value, then performed a t-test to identify parks that consistently scored higher or lower than expectations. Thus, we tested which, if any, of these well-sampled, relatively small parks differed consistently from expectations in their observed, site-level BirdsPlus scores (i.e., if the residuals of the site-specific checklists consistently differed from zero).

## RESULTS

### Acoustic checklists

For both BirdNET and Merlin, the number of species reported on each recording decreased as the detection threshold increased (Table 1). Approximately the same number of species on average were detected with the custom (species-specific) thresholds for Merlin as when a flat 0.9 threshold was used (although these were not necessarily the same species). Notably, for all models and thresholds, detections are not guaranteed to be true positives, and false positives increase as detection thresholds decrease. More species were detected when using Merlin than BirdNET (Table 1), and at a flat threshold of 0.9, 29% of recordings had no species detected with BirdNET. This had important implications for our downstream modeling power.

**Table 1.** Mean and standard deviation (SD) in species richness (SR) per checklist across detection modalities and thresholds. Mean SR per acoustic checklist decreases with increasing threshold, while the number of zeros—checklists with no species detected at all—increases.

| Modality | Mean SR | SD SR | Length of Zeros |
| --- | --- | --- | --- |
| Merlin 0.25 | 6.45 | 4.91 | 30 |
| Merlin 0.5 | 4.78 | 3.52 | 129 |
| Merlin 0.9 | 3 | 2.18 | 681 |
| Merlin custom | 3.68 | 2.74 | 351 |
| BirdNET 0.25 | 2.27 | 1.81 | 856 |
| BirdNET 0.5 | 1.49 | 1.08 | 2299 |
| BirdNET 0.9 | 0.82 | 0.64 | 8250 |
| eBird | 23.7 | 12.96 | 0 |

### Spatial biodiversity expectations

Maps of predicted total BirdsPlus species scores—the sum of all detected species’ BirdsPlus scores expected after a period of sampling (be it acoustic or eBird observer)—varied substantially between modalities and between sound ID models (Fig. 4). All of the Merlin-derived maps appeared fairly similar, with high expected diversity in the Adirondacks, Alleghenies, and Virginia Piedmont. The BirdNET-derived maps varied across the threshold used. At 0.25, the agriculture-dominated regions of western Ohio and the coastal and littoral habitats of the Chesapeake Bay, Long Island, and Lake Erie stand out for high expected diversity, whereas at 0.5 and 0.9, the expected diversity in these agriculture-dominated regions diminishes substantially. The eBird map differs from both the BirdNET and Merlin maps, with very high predicted diversity in the Chesapeake Bay, Virginia Piedmont, and littoral and riparian habitats such as those along the southern shores of Lakes Ontario and Erie, and along the St. Lawrence River. Across all modalities and thresholds, the lower diversity expected in urban areas was one of the most obvious visual patterns.

**Figure 4.**
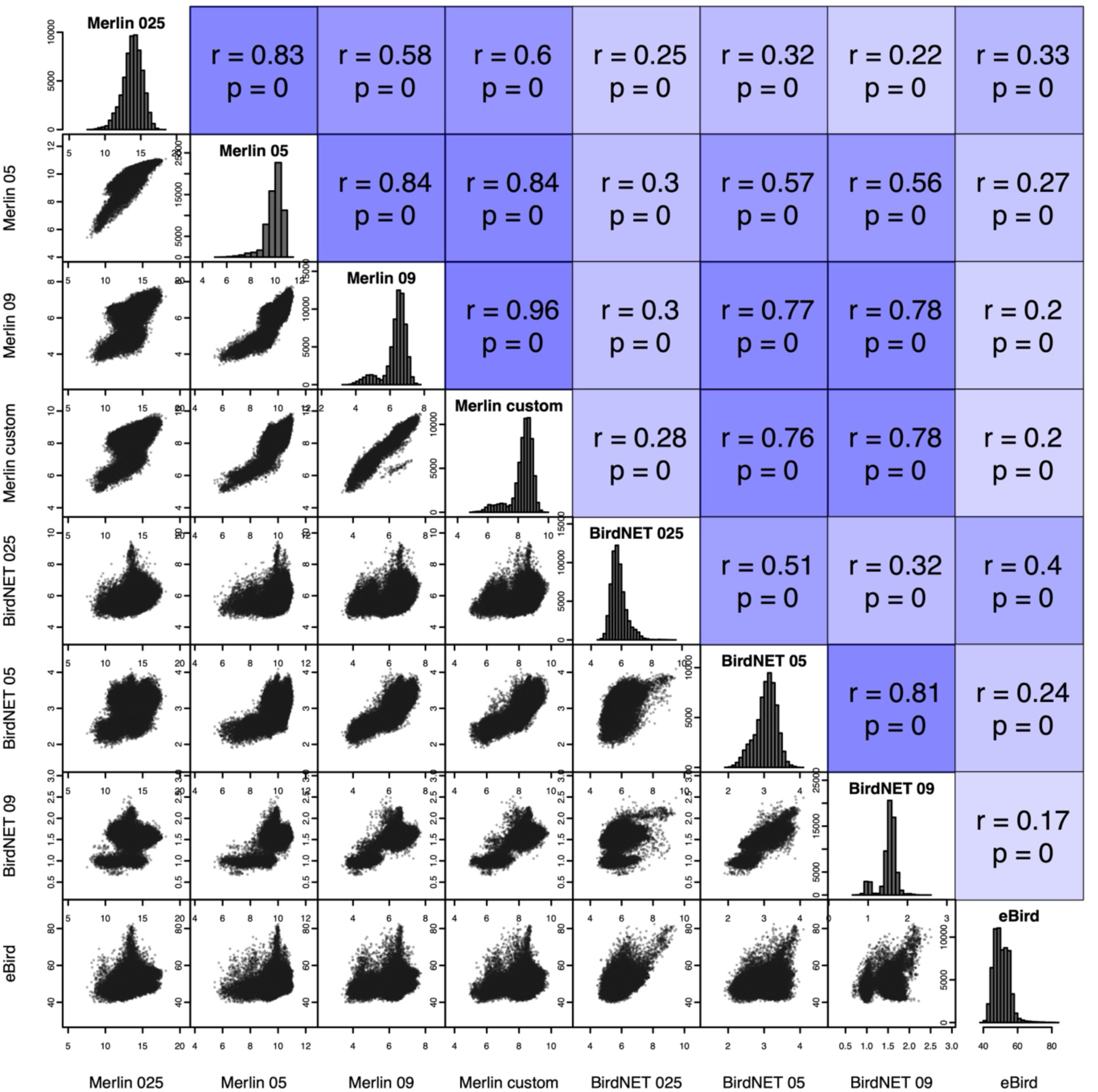
Paneled correlation plot showing the pixel-level correspondence between maps of expected total BirdsPlus scores for the different detection modalities and thresholds in the study. The Merlin maps are more similar to one another than are the BirdNET maps to each other (cluster of blue squares in the top left of the figure), but the map that aligns most closely with the eBird map is BirdNET 0.25.

The correlation strength amongst the four Merlin maps was stronger than it was amongst the BirdNET maps. Although the BirdNET 0.25 map was quite dissimilar from the other BirdNET maps, it did show a strikingly similar signal to the eBird map with respect to high expected diversity along the southern shore of Lake Erie and along the Delmarva Peninsula (Fig. 2).

### Model testing

According to basic test statistics (Table 2), performance increased at lower thresholds for both sound ID models and, across all detection thresholds, the Merlin models nominally outperformed the BirdNET models. Note, however, that these values quantify how well the fitted random forest models are able to predict BirdsPlus site scores as measured by the aggregated output of the sound ID models; they do not quantify performance based on a validated dataset of known diversity measures.

**Table 2.** Mean squared error (MSE), root mean squared error (RMSE), mean absolute error (MAE), and R^2^ for the fitted random forest models across detection modalities and thresholds. These models predict total site-level BirdsPlus scores, and are assessed against a held-out test dataset, which is itself derived from unverified detections. False positives and negatives can and do occur across modalities, and more false positive (and fewer false negative) errors occur when deriving acoustic checklists using lower thresholds.

| Modality | MSE | RMSE | MAE | R<sup>2</sup> |
| --- | --- | --- | --- | --- |
| Merlin 0.25 | 9.69 | 3.11 | 2.22 | 0.54 |
| Merlin 0.5 | 5.43 | 2.33 | 1.68 | 0.49 |
| Merlin 0.9 | 2.79 | 1.67 | 1.23 | 0.37 |
| Merlin<br>custom | 5.18 | 2.28 | 1.61 | 0.42 |
| BirdNET<br>0.25 | 4.04 | 2.01 | 1.38 | 0.30 |
| BirdNET<br>0.5 | 1.91 | 1.38 | 0.96 | 0.19 |
| BirdNET<br>0.9 | 1.11 | 1.05 | 0.80 | 0.10 |
| eBird | 48.34 | 6.95 | 5.32 | 0.74 |

Across all four Merlin-based models, the top five most important covariates were always the same: checklist duration, day of year, time of day, time relative to solar noon, and local road density (Table S1). The same was similar for the BirdNET-based models, where six covariates were always top-ranked: checklist duration, day of year, time of day, time relative to solar noon, local road density, and nighttime light density. For the eBird observer-based model, a number of sampling effort covariates also consistently ranked as most important, and across all model types, partial dependence plots corroborated expected relationships of these effort variables and predicted total BirdsPlus score—increased sampling effort resulted in an increase in detected diversity (Fig. S1). However, when focusing only on environmental variables, clear differences emerged in what the models relied on for predicting expected BirdsPlus site scores (Table S1). For example, the single-most important environmental variable for the eBird map was the percentage of seasonal water cover, as opposed to the urban-density and elevation variables that were important for the acoustic-based models. The SHAP plots combine predictor importance and partial dependences into a more intuitive, visual representations of these results (Fig. S2).

### BirdsPlus Index results

Despite the BirdNET and Merlin maps differing substantially at many detection thresholds, BPI values for the held-out parks tended to align across sound identification models. That is, the residuals of observed versus expected were at least broadly similar irrespective of the sound identification model used (Table S2, Fig. 5). This was particularly true for expectations across detection thresholds within the Merlin model, but it was also true when comparing expectations from the BirdNET model using thresholds of 0.25 and 0.5 with those from the Merlin model (Fig. 5). Despite better correspondence between the eBird observer and Merlin-based maps than the BirdNET-based maps, BPI scores derived with the acoustic models showed stronger correlation to each other than they did to eBird-based BPI scores. Still, eBird-based BPI scores did show some alignment with acoustic-based scores, particularly those derived with Merlin (e.g., the Pearson correlation coefficient between mean, site-level eBird BPI scores and Merlin 0.9 threshold BPI scores was 0.36, p = 0.05).

**Figure 5.**
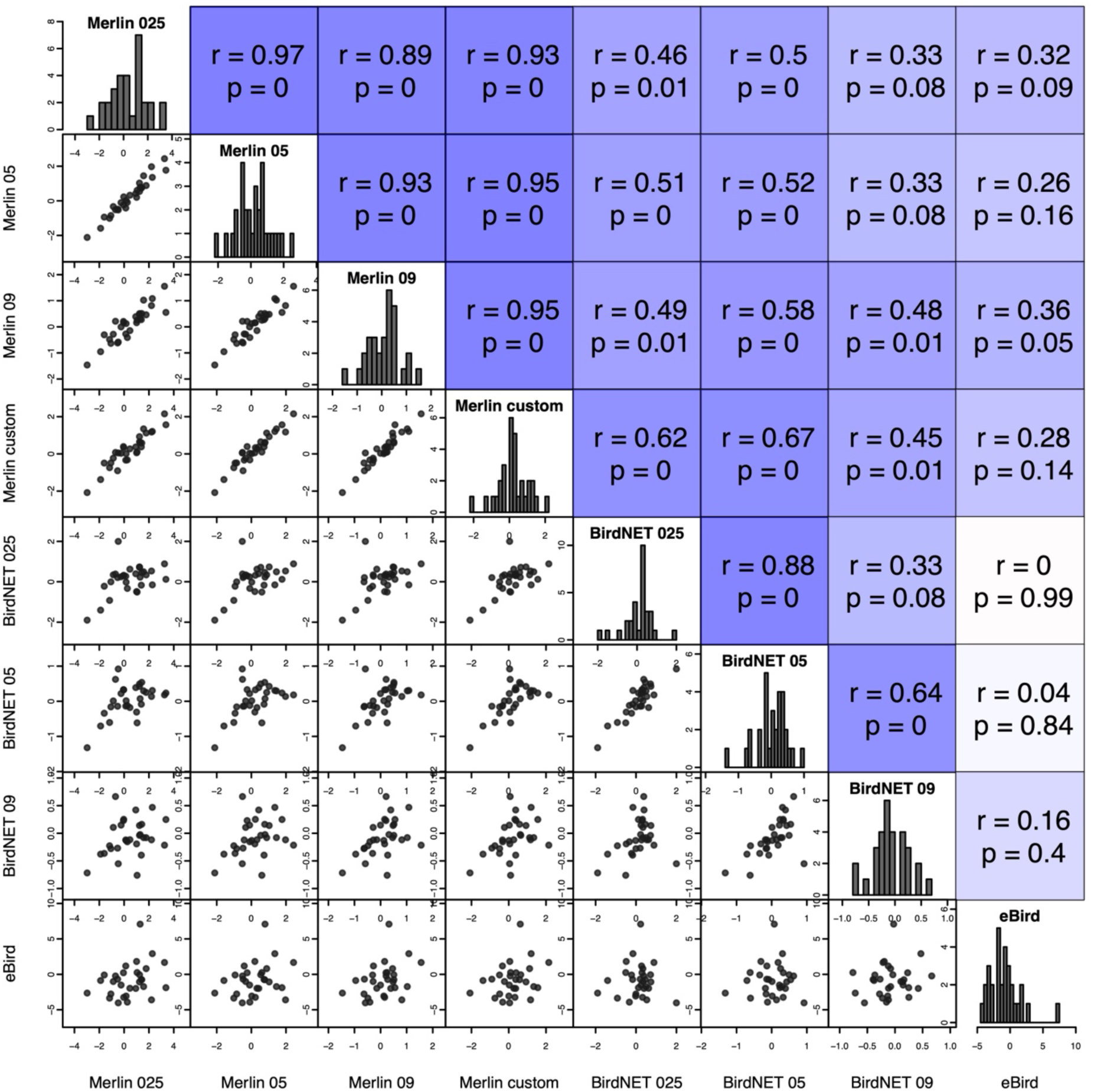
Paneled correlation plot showing the correspondence in mean BirdsPlus Index scores per focal park in the study highlights the generally divergent BPI scores when eBird observer checklists are used instead of acoustic checklists. Despite the maps of expected total site scores diverging notably across alternative sound identification models, the residuals of observed versus expected were at least broadly similar irrespective of the model used.

## DISCUSSION

In this paper we proposed a methodology (Fig. 1) for linking automated acoustic monitoring with remote sensing data to predict variation in regional biodiversity (Fig. 2). We hypothesized that this approach could be leveraged to develop a site-based index describing deviation from expectation, which we call the BirdsPlus Index (BPI). We tested this in the Northeast United States, with a particular focus on small parks. We found broad support for our hypothesis: expected conservation value aligned with known environmental drivers such as urban density and elevation (Table S1), and BPI scores from small, well-sampled parks were consistent across a suite of automated acoustic identifiers and thresholds (Table S2, Fig. 5). While BPI scores showed less correspondence with expectations derived from eBird observer checklists^24^, our results nonetheless establish both a novel conservation value metric—globally scalable and likely to improve in accuracy over time—and the foundation for a general framework for integrating acoustic data into global biodiversity monitoring.

One of the most appealing aspects of our methodology is its accessibility. In principle, anyone with a smartphone could assess habitat quality in real time by using their device to listen to the world around them. While this remains a dream in many regions of the globe, our results point towards a path forward where data coverage allows. Importantly, this vision calls for acoustic sampling that is broad across time and space, but not necessarily intensive at any single location. Current passive acoustic monitoring projects often sample intensively at limited sites, generating vast sums of recordings—for example, a recent project of our own yielded over 40,000 hours of recordings from only two small farms in Costa Rica. Many such efforts aim to compile complete species lists, but until automated classifiers approach perfection, this goal requires manual validation at often infeasible scales^48^. Moreover, complete species lists may not be the precise goal we should pursue. Reconsidering acoustic monitoring as a sampling strategy with imperfect detection and measurable false positive rates is a compelling alternative.

Our approach avoids the problem of exhaustive manual validation by passing a spatiotemporally broad sample of recordings through a single classifier, deriving expectations, and comparing focal recordings to those expectations (Fig. 1). This design assumes classifier accuracy is relatively constant over time and space—an assumption that is imperfect but reasonable for our proof of concept. Indeed, we obtained similar BPI scores for acoustic-based models across a range of detection thresholds, even with an extremely limited dataset (e.g., just 324 hours of acoustic training data compared to 17,561 hours for eBird, Fig. 5). With an average of only 30 minutes of sampling per park, we still derived site-level BPI scores that differed consistently from expectations (Table S2). Nevertheless, many projects will continue to require sustained high-intensity monitoring and manual human validation, such as surveys for threatened^49^ or lost species^50^, or habitat improvement projects that rely on indicator species as measures of success^9^.

The greatest current limitation of our approach is species coverage in sound identification models. In the Northeast US, 82% of species reported on paired eBird checklists are covered by Merlin (v43). The majority of those not covered would not be detected acoustically even if they were included in the classifier, as they either vocalize extremely infrequently (e.g., Turkey Vulture, *Cathartes aura*), or are vagrants to the region and would have been removed by our spatiotemporal filtering process. In contrast, species coverage drops to ∼25% in Peru and ∼10% in Australia, and many of the currently unincluded species in these regions could theoretically be detected by their vocalizations. These coverage gaps are not random: species that are rare, vocalize infrequently, or are from rarely accessed habitats are often poorly represented in sound libraries (and therefore are also absent from annotated datasets). This asymmetry risks skewing results. For instance, in comparing shade-grown versus sun-grown coffee plantations, current models might suggest higher biodiversity in the latter simply because forest specialists are underrepresented in sound identification models. Targeted sound recording to expand classifier coverage should be a global priority.

Our results also highlight the potential for a lack of recorder diversity to bias results, and a need for complementary acoustic metadata to help mitigate this potential bias. In our focal sites, parks with high BPI scores tended to have diverse habitats and recordings from multiple contributors, while low-BPI sites were often dominated by single recordists using directional microphones. Such microphones are able to minimize captured background “noise”, and a key goal of those using such equipment is to produce clear and long recordings with little audible aside from the target species. These recordings generally sound better, and they make excellent material for teaching, outreach, and training sound ID models (image augmentation techniques can be used to simulate overlapping vocalizations). Yet, in our framework the background species—the “noise” so assiduously avoided in high-quality recordings—are critical data, and the slow accumulation of species over long-duration recordings results in a low BPI score (Fig. S1). Acoustic metadata describing recording equipment would help mitigate this issue^51^. Similarly, recordings of particularly poor quality (where birds are barely distinguishable from background) also led to low BPI scores. These patterns suggest our approach benefits from diverse contributors and would improve with larger, more heterogeneous datasets.

Differences between eBird observations and acoustic monitoring are largely beyond the scope of the current manuscript, but we can identify a few possible reasons that results did not perfectly align. Our study period, June through August, was an attempt to optimize the number of recordings we could include in the model while ideally restricting our analysis to the breeding season. In practice, particularly across this broad of a latitudinal range, we are capturing migrants heading northward at the start of the study period, and southward towards the end of it^27^. While this may not overly affect the resulting acoustic-based maps, since migrants tend to vocalize less often, it certainly affected the eBird observer maps—sites with high expected total diversity for the eBird-based approach were often coastal and littoral sites, and the highest diversity checklists for these sites tended to be in August. This is when shorebirds (and many other species) are migrating southward, and observers station themselves along shorelines to monitor their passage. Not only do many of these species vocalize infrequently (and briefly, from the perspective of an observer on the ground), but many of these shorebirds have high BirdsPlus species-scores^34^. The result was that, particularly given the criteria we chose for data inclusion, the maps of expected diversity did not closely align between detection modalities (Figs. 3-4). Nevertheless, these expectations partly mitigated differences when comparing the residuals of the site-level observations against expectations, such that BPI scores showed some alignment between eBird and acoustic-based models (Fig. 5). We suspect that if we had access to more acoustic data, and could restrict our study to a thinner slice of time (e.g. mid-June through mid-July), the models would show further alignment. That said, these are fundamentally different detection methods— acoustic monitoring should not be viewed as an alternative to surveys by experienced observers—and we should not expect these maps, and the resulting BPI scores, to perfectly match.

We propose several solutions to tackle current challenges associated with widespread acoustic monitoring, and with the uptake of such methodologies to inform metrics of ecological integrity. The most direct path to improving classifier coverage is citizen science. Just as eBird^24,25,52^ and iNaturalist^53^ have closed major monitoring gaps through mass participation, initiatives that encourage sound recording can help expand sound libraries, which ultimately form the basis of annotated data to improve classifiers^22^. Strategic deployment of recording devices also matters: our results suggest that broad, distributed, short-duration recordings may be more useful for biodiversity and conservation value assessment than intensive sampling at a few sites. Integration of these methodologies with global sustainability frameworks is another key next step. Metrics like the BPI could support initiatives such as the Science Based Targets Network^54^, Verra’s Nature Framework^55^, and Plan Vivo^56^, where intuitive conservation value and outcomes measures grounded in real biodiversity monitoring data are urgently needed. Acoustic monitoring should be viewed not as a replacement for traditional surveys, but as a complementary tool that extends coverage, provides a long-term record that can be revisited over time, and that incorporates measurable error rates^16,17^.

Our motivation in developing the BPI is to provide a robust, intuitive biodiversity metric that captures the benefits of sustainable management practices such as managed timber forests^57^, shade-grown coffee, and reforestation programs^58^. More broadly, our approach lays the groundwork for a global acoustic biodiversity monitoring framework. As sound libraries continue to expand and classifiers improve, this framework could support real-time, large-scale conservation assessments while creating a lasting archive of global soundscapes. Without such a record, our ability to compare present-day ecosystems to past conditions will remain limited. But if built now, a time-stamped global library of recordings will allow us to readily quantify shifts in biodiversity and conservation outcomes as they unfold over the coming decades.

## ACKNOWLEDGMENTS

We thank Daniel Fink, Tom Auer, Courtney Davis, Viviana Ruiz-Gutierrez, and Matthew Strimas-Mackey for advice and consultation with distribution modeling and mapping. We thank David Hartwell for initially encouraging us to pursue an avian, acoustic-based monitoring tool that resulted in the development of this methodology.

## SUPPLEMENTARY MATERIALS

Table S1. Mean predictor importance across detection modalities and thresholds, sorted by decreasing average importance across all models.

Table S2. Park name, size (in square meters), mean latitude and longitude of the park-specific acoustic and observer checklists that went into informing the acoustic and eBird-observer-based BirdsPlus Index scores shown here.

**Figure S1.**
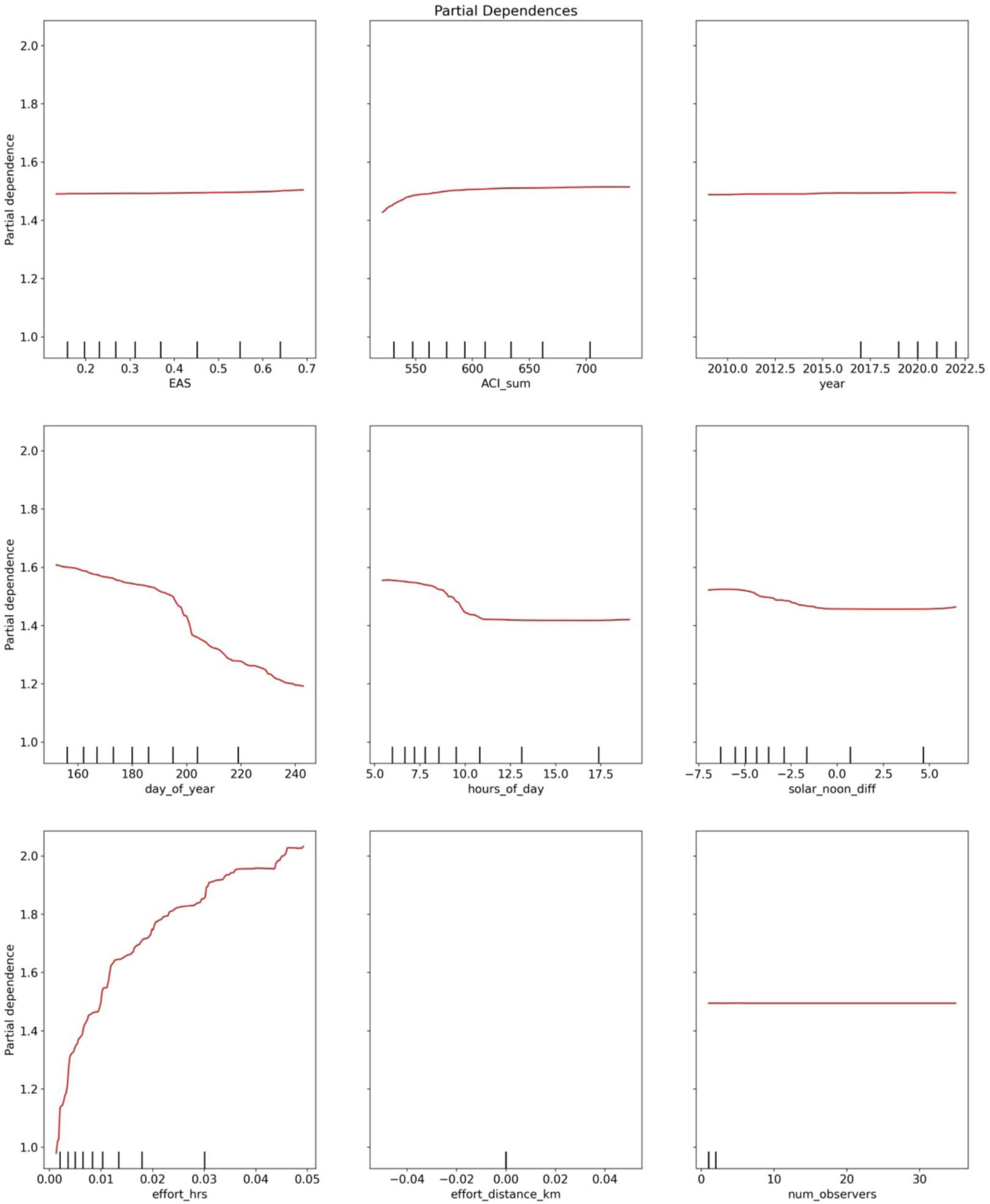
We employed partial dependence plots to assess model performance and identify, for mapping purposes, the values of effort variables that maximized detection. The example shown is the partial dependence plot for the Merlin-based model with a custom (species-specific) detection threshold.

**Figure S2.**
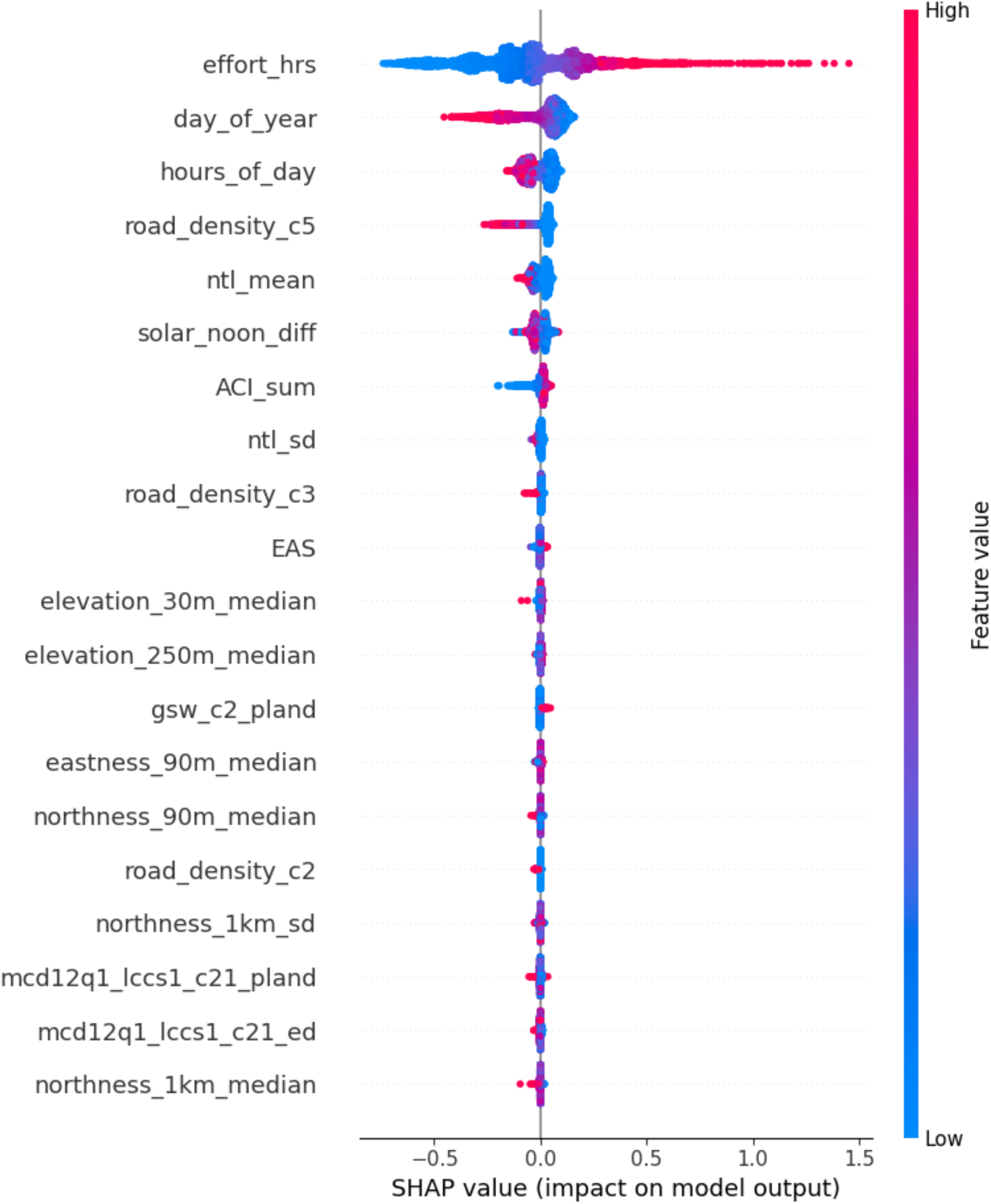
We employed SHAP plots to assess model performance and gauge the importance and directionality of the many covariates used to model expected BirdsPlus site scores (expected total biodiversity). The example shown is for the Merlin-based model with a custom (species-specific) detection threshold. Descriptions of these covariates are found in Table S1.

